# Exploring building blocks of cell organization by estimating network motifs using graph isomorphism network

**DOI:** 10.1101/2023.11.04.565623

**Authors:** Yang Yu, Shuang Wang, Dong Xu, Juexin Wang

## Abstract

The spatial arrangement of cells within tissues plays a pivotal role in shaping tissue functions. A critical spatial pattern is network motif as the building blocks of cell organization. Network motifs can be represented as recurring significant interconnections of cells with various types in a spatial cell-relation graph, i.e., enriched occurrences of isomorphic subgraphs in the graph, which is computationally infeasible to have an optimal solution with large-size (*>*3 nodes) subgraphs. We introduce <u>Tri</u>angulation Network <u>M</u>otif <u>N</u>eural <u>N</u>etwork (TrimNN), a neural network-based approach designed to estimate the prevalence of network motifs of any size in a triangulated cell graph. TrimNN simplifies the intricate task of occurrence regression by decomposing it into several binary present/absent predictions on small graphs. TrimNN is trained using representative pairs of predefined subgraphs and triangulated cell graphs to estimate overrepresented network motifs. On typical spatial omics samples within thousands of cells in dozens of cell types, TrimNN robustly infers the presence of a large-size network motif in seconds. In a case study using STARmap Plus technologies, TrimNN identified several biological meaningful large-size network motifs significantly enriched in a mouse model of Alzheimer’s disease at different months of age. TrimNN provides an accurate, efficient, and robust approach for quantifying network motifs, which helps pave the way to disclose the biological mechanisms underlying cell organization in multicellular differentiation, development, and disease progression.

## 1 Background

Deciphering the relationship between structure and function in tissues is the cornerstone of tissue biology and pathology[15]. Unraveling the mechanisms underlying the spatial organization of different cell types in a specific tissue is pivotal for addressing this fundamental biological question. With advancements in spatial omics, such as spatially resolved transcriptomics[18] and proteomics[11], researchers have access to unprecedented resources to explore how distinct cell types are organized to perform specialized roles at the cellular level[3]. Typically, the spatial cell organization obtained through spatial omics is predominantly focus on assessing the cell type composition. However, identifying the building blocks of the cell organization and determining which spatial cellular interconnection patterns are informative to tissue function remains challenging[2].

Network motifs as recurring significant interconnections represent network characteristics as conservative patterns. The concept of network motif was introduced by Uri Alon in 2002[14], and subsequent studies have greatly enhanced the knowledge of network functions in social networks[22] and biological networks[16], such as metabolic networks[1], gene-regulatory networks[20], and protein-protein interaction networks[29]. We hypothesize network motifs can be treated as building blocks of cell organization that invariantly across different samples, and they connect with key functions in a biologically meaningful context. Compared to classical network motifs, such as the feed-forward loop (FFL)[12], cell organization-associated network motifs do not have directions but are instead characterized by node labels representing distinct cell types. Typically, a network motif analysis involves counting the occurrences of various combinations of cell types, including size-1 motifs (nodes), size-2 motifs (nodes on edges), size-3 motifs (triangles), and large-size motifs (more nodes than a triangle). Currently, most existing network motif analyses are limited to size 1-3[27]. However, in spatial omics studies, biologists have observed the prevalence of large-size network motifs significantly correlated with patient survival in colorectal cancer[19], brain tumor[7], and lung cancer[21].

This biological problem of identifying overrepresented network motifs can be modeled mathematically by identifying the most overrepresented subgraphs. This problem usually consists of two sub-problems: subgraph matching[23] and pattern growth[16]. Given a specific subgraph, subgraph matching counts the occurrence of isomorphic subgraphs within the overall graph. Building upon subgraph matching, the pattern growth problem enumerates and identifies the most overrepresented subgraphs based on their occurrences in the subgraph matching. It is proven that subgraph matching is NP-complete[5], which makes it computationally infeasible to count large-size isomorphic subgraphs in polynomial time. Even though many methods adopted heuristic strategies, such as edge sampling (e.g., MFinder[8]), node sampling (e.g., FANMOD[26]), and global pruning (e.g., Ullmann[24] and VF2[4]) to address this challenge, their practical utility remains limited due to the scalability issue. Neural Subgraph Isomorphism Counting (NSIC)[10] is the first deep learning model using graph neural networks to predict subgraph occurrences as a regression problem, but its far-reaching goal on universal graphs and its limited accuracy narrow its practical utility.

Here, we present <u>Tri</u>angulation Network <u>M</u>otif <u>N</u>eural <u>N</u>etwork (TrimNN), a neural network-based approach to estimate the prevalence of network motifs of any size in a graph. TrimNN aims to address the subgraph matching problem in triangulated graphs derived from spatial omics. TrimNN decomposes the occurrence regression challenge into several binary classification problems modeled by the sub-TrimNN module. Inspired by NSIC, TrimNN is trained on representative pairs of the predefined subgraphs and the triangulated cell graphs. TrimNN aggregates the sub-TrimNN module’s results and outputs the subgraphs’ relative abundance, which can be used to estimate the prevalence of overrepresented network motifs.

Our major contribution is formulating and simplifying the subgraph matching challenge from a regression problem to many binary classification problems in the context of spatial omics. Avoiding predicting the absolute occurrences of network motifs in the universal graphs in the original setting, TrimNN decomposes the challenge to a serial binary classification problem in a well-defined set of biological meaningful triangulated graphs, where it performs binary presence/absence predictions at a similar scale. TrimNN only models the specific triangulated graphs after Delaunay triangulation on spatial omics data, where the spatial space is filled with only triangles. Most importantly, TrimNN can be used to output the most overrepresented network motifs with top relative abundances, simplifying the problem and bypassing the challenging occurrence counting. TrimNN is publicly available at https://github.com/yuyang-0825/TrimNN.

## 2 Methods

### 2.1 Problem setting and definition

Formally, we define the triangulated graph G from spatial omics as *G* = {*V, E*}. The size-*k* subgraphs with *k* nodes as *M*_*k*_ are induced subgraphs of *G*. The biological problem of identifying the overrepresented network motifs can be modeled mathematically in finding the most overrepresented subgraphs 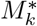 in *G*, where 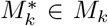 and *M*_*k*_ ∈ *G*. This challenge consists of a subgraph matching problem and a pattern growth problem built on it. The whole workflow is shown in Figure 1.

**Fig. 1.**
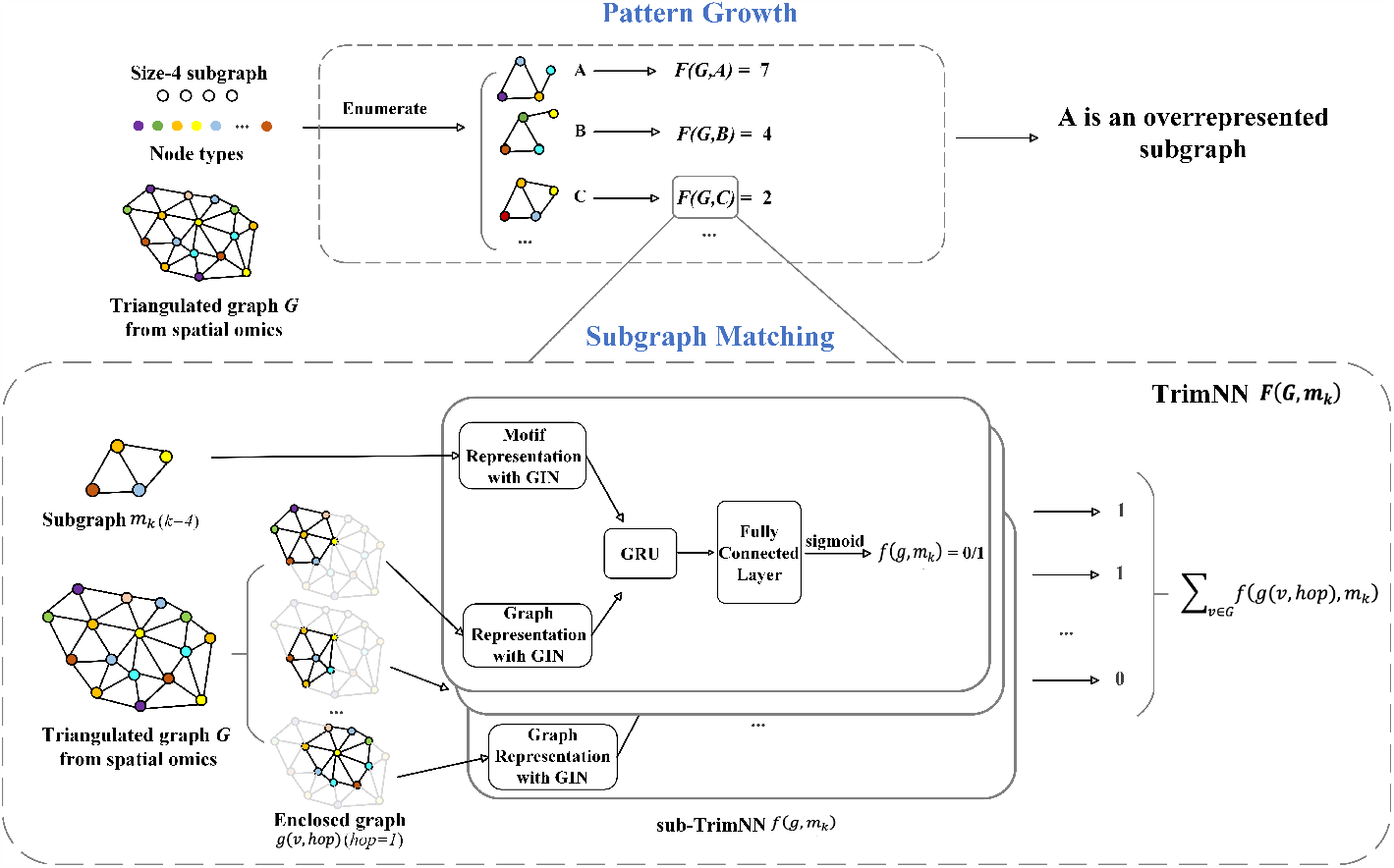
Flowchart of the motif identification problem and the TrimNN framework. The whole process comprises Pattern Growth (top) and Subgraph Matching (bottom). A size-4 subgraph *m*_*k*_ is taken as an example. In Pattern Growth, all possible size-4 subgraphs are enumerated in a triangulated graph *G* with trained TrimNN function *F* (*G, m*_*k*_) from Subgraph Matching. TrimNN summarizes many sub-TrimNN functions *f* (*g, m*_*k*_), which make binary predictions on presence (1) or absence (0) in all possible enclosed graph *g* within *G*.

The goal of TrimNN is the subgraph matching problem, which aims to define *F* (*G, m*_*k*_) ∈*N*, estimating the relative occurrence of the given *m*_*k*_ in *G*. The problem can be quasi-divided and conquered by summarization of many sub-TrimNN problems. The goal of sub-TrimNN is to build a reliable binary prediction model *f* (*g, m*_*k*_) ∈ [0, 1], where 0 presents *m*_*k*_ is absent in *g*, and 1 represents presence. Here *g ∈ G* and *g* is in a similar scale of *m*_*k*_. With sub-TrimNN on enclosed graphs of each node, TrimNN is the summarization of results from all sub-TrimNN in the graph. Finally, *F* (*G, m*_*k*_) = Σ_*v*∈*G*_ *f* (*g*(*v, hop*), *m*_*k*_), where *g*(*v, hop*) is the enclosed graph as the neighborhoods of node *v* ∈ *V* with *hop* ∈ [1, 2, 3, …], and *g*(*v, hop*) ∈ *G*. The value of *hop* is related with the length of the longest path of *m*_*k*_. After we get a fast and reliable *F* (*G, m*_*k*_) from TrimNN, we can use it in the problem of pattern growth. Using enumeration or other searching processes, the final target top overrepresented set 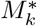 has the maximum relative abundance identified by *F* (*G, m*_*k*_), where 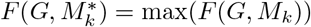.

### 2.2 TrimNN Model Architecture

We decompose the regression problem of TrimNN in the whole triangled graph into many binary classification problems in small, enclosed graphs solved by sub-TrimNN. The input of sub-TrimNN is a pair of subgraph *m*_*k*_ and the triangulated graph *g*, both of which can be extracted and learned by Graph Isomorphism Network (GIN)[28]. Then this pair of representations are aligned in the interaction module with gated recurrent units (GRU). After fully connected layers and activated by the sigmoid function, sub-TrimNN outputs the binary predictions (presence/absence). The training process minimizes the loss function on cross-entropy of known presence/absence relations. After trained sub-TrimNN *f* (*g, m*_*k*_), TrimNN estimates the abundances of *F* (*G, m*_*k*_) by summarizing sub-TrimNN predictions on each node’s enclosed graph.

### 2.3 Constructing the training set as pairs between subgraphs and triangulated graphs

In spatial omics samples, the spatial relations between the cells can be modeled as a cell graph by their coordinates in the region using the Delaunay triangulation[19]. In the triangulated graph, each node denotes a cell and is labeled with a cell type, and each edge represents the hypothetical spatial relations between cells.

We simulated the training set on known presence/absence relations of pairs between subgraphs and triangulated graphs. 8 distinct subgraphs in various topologies were generated, including size-3 and large-size subgraphs up to size-9. Given the context of routine spatial omics for each network motif, we constructed the corresponding triangulated graphs with varying node sizes of 16, 32, 64, and 128, and node types of 8, 16, and 32. Each pair of subgraph and triangulated graph has the same number of node types. To preserve the diversity, we generated 50 extended subgraphs permutating node types for each subgraph and corresponding 1,000 triangulated graphs permutating node types. We controlled the proportion of positive to negative samples at 1:1 in data generation and split the generated data into training, validation, and test sets in a ratio of 8:1:1.

## 3 Results

We compared the performance of TrimNN with competitive representative enumerating searching method VF2 and neural network based NSIC.

### 3.1 TrimNN accurately identifies the presence of the network motifs in a triangulated graph

We first tested the performance of TrimNN on a modified task of subgraph matching, which predicts whether the network motif existed in the triangulated graph by sub-TrimNN. As a binary prediction task, the performance was evaluated by precision, recall, F1 score, and MCC (Matthews Correlation Coefficient) on the generated test set of varying size and node types. For NSIC regresses continuous occurrence, we treat NSIC’s prediction on 0 as not existing in the graph, and any value equal to or larger than one as existing in the graph. We selected four distinct network motifs for testing, one in size-3, two in size-4, and one in size-5 (Table 1). We did not include VF2 in the performance comparison but used VF2 to generate the ground truth by enumerating all the possibilities and guaranteeing the exact results with a huge computational cost. We observed that TrimNN outperforms the competitor’s results in nearly all the scenarios (Table 1). Notably, we value the near-perfect performance on recall criteria, indicating TrimNN is confident when it predicts the network motif presented in the triangulated graph.

**Table 1.**
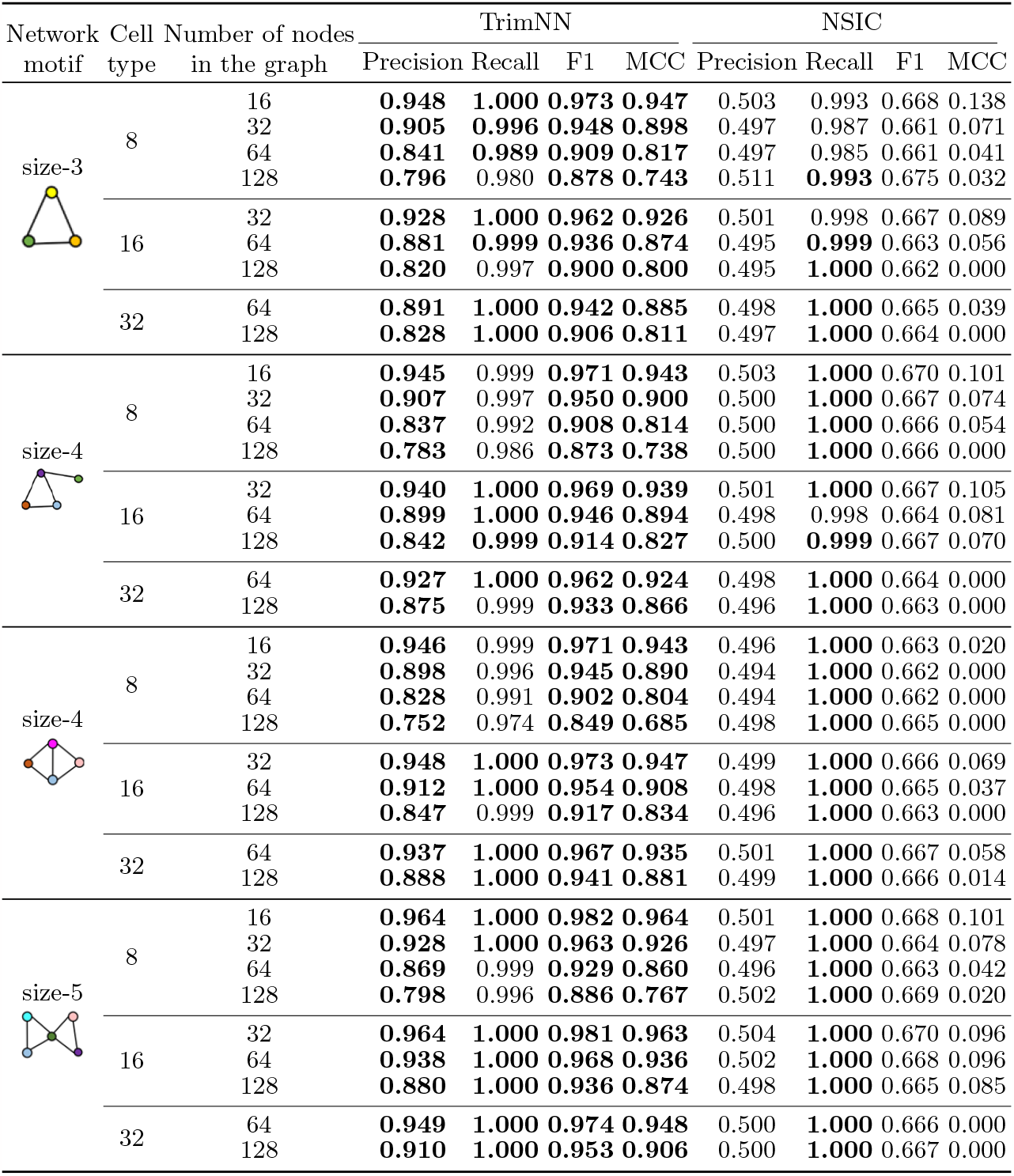
The performance of TrimNN and NSIC.

| Network motif | Cell type | Number of nodes in the graph | TrimNN |  |  |  | NSIC |  |  |  |
| --- | --- | --- | --- | --- | --- | --- | --- | --- | --- | --- |
|  |  |  | Precision | Recall | F1 | MCC | Precision | Recall | F1 | MCC |
| size-3 | 8 | 16 | **0.948** | **1.000** | **0.973** | **0.947** | 0.503 | 0.993 | 0.668 | 0.138 |
|  |  | 32 | **0.905** | **0.996** | **0.948** | **0.898** | 0.497 | 0.987 | 0.661 | 0.071 |
|  |  | 64 | **0.841** | **0.989** | **0.909** | **0.817** | 0.497 | 0.985 | 0.661 | 0.041 |
|  |  | 128 | **0.796** | 0.980 | **0.878** | **0.743** | 0.511 | **0.993** | 0.675 | 0.032 |
|  | 16 | 32 | **0.928** | **1.000** | **0.962** | **0.926** | 0.501 | 0.998 | 0.667 | 0.089 |
|  |  | 64 | **0.881** | **0.999** | **0.936** | **0.874** | 0.495 | **0.999** | 0.663 | 0.056 |
|  |  | 128 | **0.820** | 0.997 | **0.900** | **0.800** | 0.495 | **1.000** | 0.662 | 0.000 |
|  | 32 | 64 | **0.891** | **1.000** | **0.942** | **0.885** | 0.498 | **1.000** | 0.665 | 0.039 |
|  |  | 128 | **0.828** | **1.000** | **0.906** | **0.811** | 0.497 | **1.000** | 0.664 | 0.000 |
| size-4 | 8 | 16 | **0.945** | 0.999 | **0.971** | **0.943** | 0.503 | **1.000** | 0.670 | 0.101 |
|  |  | 32 | **0.907** | 0.997 | **0.950** | **0.900** | 0.500 | **1.000** | 0.667 | 0.074 |
|  |  | 64 | **0.837** | 0.992 | **0.908** | **0.814** | 0.500 | **1.000** | 0.666 | 0.054 |
|  |  | 128 | **0.783** | 0.986 | **0.873** | **0.738** | 0.500 | **1.000** | 0.666 | 0.000 |
|  | 16 | 32 | **0.940** | **1.000** | **0.969** | **0.939** | 0.501 | **1.000** | 0.667 | 0.105 |
|  |  | 64 | **0.899** | **1.000** | **0.946** | **0.894** | 0.498 | 0.998 | 0.664 | 0.081 |
|  |  | 128 | **0.842** | **0.999** | **0.914** | **0.827** | 0.500 | **0.999** | 0.667 | 0.070 |
|  | 32 | 64 | **0.927** | **1.000** | **0.962** | **0.924** | 0.498 | **1.000** | 0.664 | 0.000 |
|  |  | 128 | **0.875** | 0.999 | **0.933** | **0.866** | 0.496 | **1.000** | 0.663 | 0.000 |
| size-4 | 8 | 16 | **0.946** | 0.999 | **0.971** | **0.943** | 0.496 | **1.000** | 0.663 | 0.020 |
|  |  | 32 | **0.898** | 0.996 | **0.945** | **0.890** | 0.494 | **1.000** | 0.662 | 0.000 |
|  |  | 64 | **0.828** | 0.991 | **0.902** | **0.804** | 0.494 | **1.000** | 0.662 | 0.000 |
|  |  | 128 | **0.752** | 0.974 | **0.849** | **0.685** | 0.498 | **1.000** | 0.665 | 0.000 |
|  | 16 | 32 | **0.948** | **1.000** | **0.973** | **0.947** | 0.499 | **1.000** | 0.666 | 0.069 |
|  |  | 64 | **0.912** | **1.000** | **0.954** | **0.908** | 0.498 | **1.000** | 0.665 | 0.037 |
|  |  | 128 | **0.847** | 0.999 | **0.917** | **0.834** | 0.496 | **1.000** | 0.663 | 0.000 |
|  | 32 | 64 | **0.937** | **1.000** | **0.967** | **0.935** | 0.501 | **1.000** | 0.667 | 0.058 |
|  |  | 128 | **0.888** | **1.000** | **0.941** | **0.881** | 0.499 | **1.000** | 0.666 | 0.014 |
| size-5 | 8 | 16 | **0.964** | **1.000** | **0.982** | **0.964** | 0.501 | **1.000** | 0.668 | 0.101 |
|  |  | 32 | **0.928** | **1.000** | **0.963** | **0.926** | 0.497 | **1.000** | 0.664 | 0.078 |
|  |  | 64 | **0.869** | 0.999 | **0.929** | **0.860** | 0.496 | **1.000** | 0.663 | 0.042 |
|  |  | 128 | **0.798** | 0.996 | **0.886** | **0.767** | 0.502 | **1.000** | 0.669 | 0.020 |
|  | 16 | 32 | **0.964** | **1.000** | **0.981** | **0.963** | 0.504 | **1.000** | 0.670 | 0.096 |
|  |  | 64 | **0.938** | **1.000** | **0.968** | **0.936** | 0.502 | **1.000** | 0.668 | 0.096 |
|  |  | 128 | **0.880** | **1.000** | **0.936** | **0.874** | 0.498 | **1.000** | 0.665 | 0.085 |
|  | 32 | 64 | **0.949** | **1.000** | **0.974** | **0.948** | 0.500 | **1.000** | 0.666 | 0.000 |
|  |  | 128 | **0.910** | **1.000** | **0.953** | **0.906** | 0.500 | **1.000** | 0.667 | 0.000 |

### 3.2 TrimNN accurately identifies top overrepresented network motifs

Then we tested whether TrimNN identifies the correct overrepresented network motifs in the triangulated graph as a pattern growth problem. To evaluate the performance, we used metrics, such as Hit Ratio (HR), Root Mean Square Error (RMSE), Mean Absolute Error (MAE), Pearson Correlation Coefficient (PCC), and Spearman Correlation Coefficient (SCC) to quantify the accuracy of the predicted occurrences. Similar to Section 3.1, we enumerated all their possibilities in both 8 cell types and 16 cell types with VF2 using significant computational resources. As we only care about the biological meaningful top overrepresented network motifs in practice, we tested whether these methods can successfully identify the top 5 and top 10 overrepresented network motifs. The overall performance in identifying the occurrences of all the possible network motifs were also evaluated. The results in comparing the occurrences of network motifs are shown in Table 2.

**Table 2.**
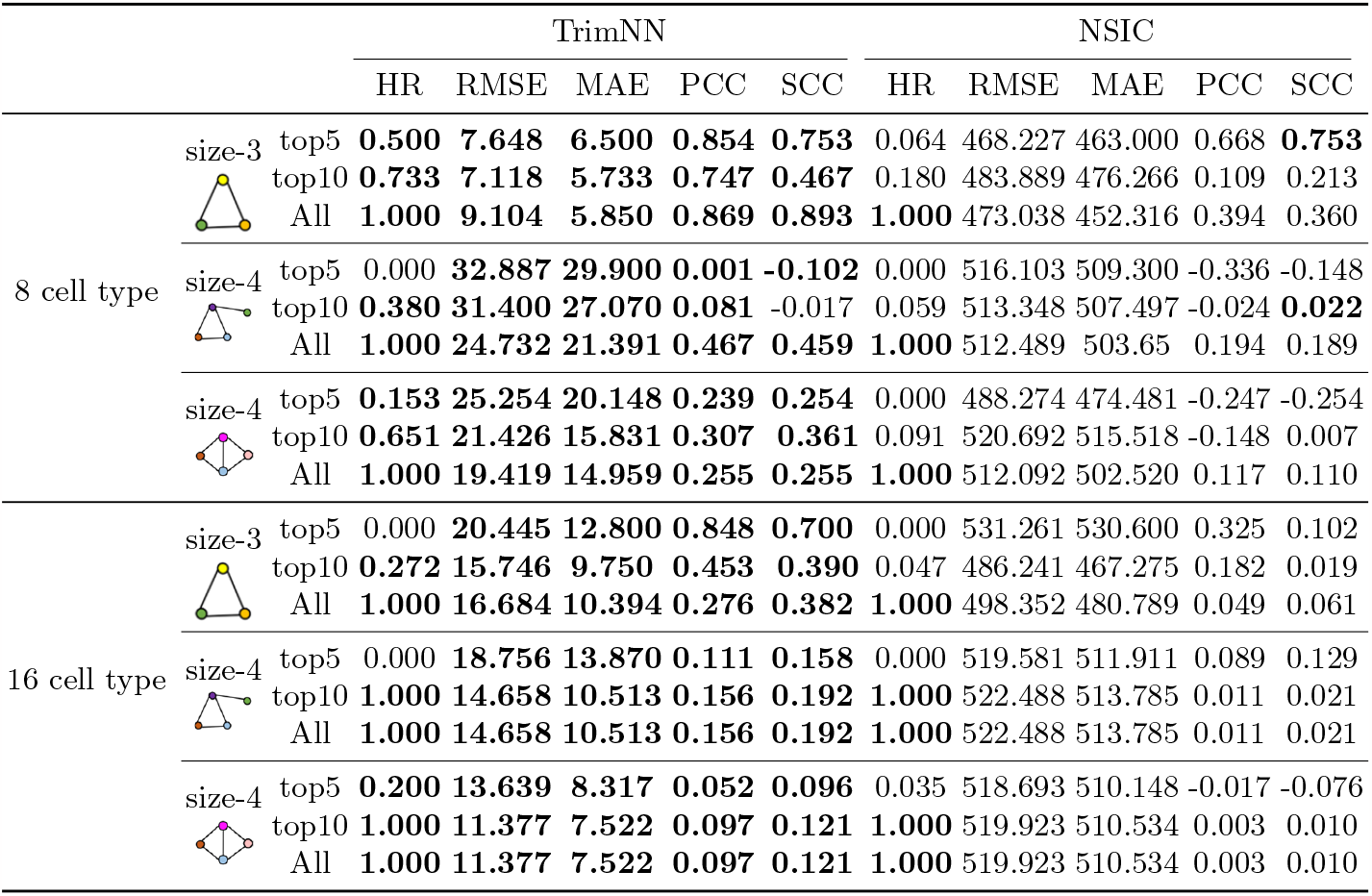
The Performance of TrimNN and NSIC in identifying the occurrences of top overrepresented network motifs.

We observe that TrimNN constantly outperforms NSIC in most scenarios and most criteria by a large margin. Notably, TrimNN demonstrates an average improvement over NSIC by approximately 50 times in RMSE and MAE. In TrimNN, we observe the performance of the second size-4 network motif is consistently better than the first size-4 network motif. This difference may result from the inductive bias of topology. If the network motif is composed of triangles, it tends to get better performance. We also observe when the number of cell types and the size of network motifs increase, accurately identifying the top overrepresented network motifs becomes even more challenging. Besides the absolute value of occurrence, we evaluated the results as the relative value of ranking in **Supplementary Table 1**. We observe the same phenomenon that TrimNN outperforms NSIC in identifying the most overrepresented network motifs. From these results, it is evident that TrimNN accurately identifies top overrepresented network motifs.

### 3.3 TrimNN is highly scalable in identifying large-size network motifs

As scalability plays a vital role in the study, we compared the computational time on triangulated graphs with varying node sizes. In the experiments, the inquiry subgraph contains 9 nodes, both the subgraph and the triangulated graph have 32 node types. All the experiments were performed on a workstation equipped with AMD EPYC 7713 CPU and one NVIDIA A100 GPU. Figure 2 shows that both TrimNN (red line) and NSIC (green line) exhibit linear scalability with increasing node sizes (black dot line), while TrimNN continuously consumes even lower computational time when the graph size is in the scale of typical spatial omics samples (*>*512 nodes). In contrast, the VF2 method (blue line) grows exponentially. When the number of graph nodes exceeds 10k, VF2’s runtime surpasses 20k seconds, making it unacceptable in most scenarios.

**Fig. 2.**
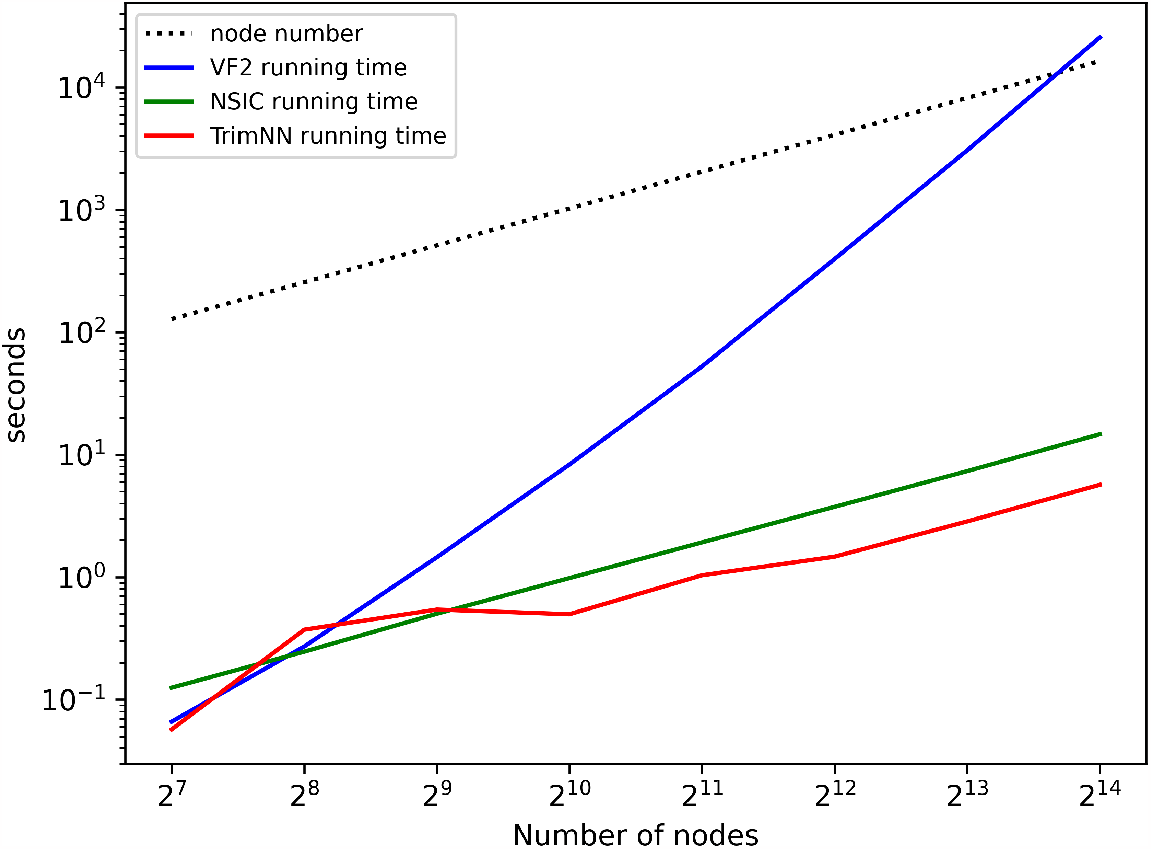
Comparison in scalability between TrimNN and competitors.

Theoretically, both the time and space complexity of sub-TrimNN inference are *O*(|*V* |+ *k*), which is linear to the node sizes of the input subgraph and the triangulated graph. The time complexity of the entire TrimNN inference is the graph node size multiples the time complexity of sub-TrimNN on all the enclosed graphs, which is *O*(|*V* | * (|*V* | + *k*)), the time complexity of building an enclosed graph is *O*(|*V* |^*hop*+1^), and the space complexity of the entire TrimNN inference is *O*(|*V*| ^*hop*^ * (|*V*| + *k*)). On the other hand, the space complexity of VF2 is *O*(*V*) and time complexity is *O*(*V* ! \**V*). In practical usage, on typical spatial omics data with thousands of cells of dozens of cell types, TrimNN robustly infers large-size network motifs accurately in seconds, which is unattainable through conventional methods.

### 3.4 Ablation tests

#### Utilized GIN is a suitable graph representation method in subgraph matching

The performance of TrimNN is heavily dependent on the expressive power of utilized GIN in representing the graph structure and features. To validate the effectiveness of GIN, we tested the model performance by replacing GIN with another two widely used graph neural network architectures, i.e., Graph Attention Networks (GAT)[25] and Graph Convolutional Networks (GCN)[9]. All other model components and parameters remained unchanged. The results are evaluated using the same criteria as Section 3.1. The results of the comparative analysis are shown in Table 3. Based on this ablation test, GIN in TrimNN consistently yields better performance than GAT and GCN, consistent with the theoretical analysis that GIN is the most powerful 1-order graph neural network[28]. This ablation test indicates that the utilized GIN is suited for capturing the characteristics of isomorphic graphs in our task, enabling the extraction of more informative graph features for subsequent subgraph matching.

**Table 3.**
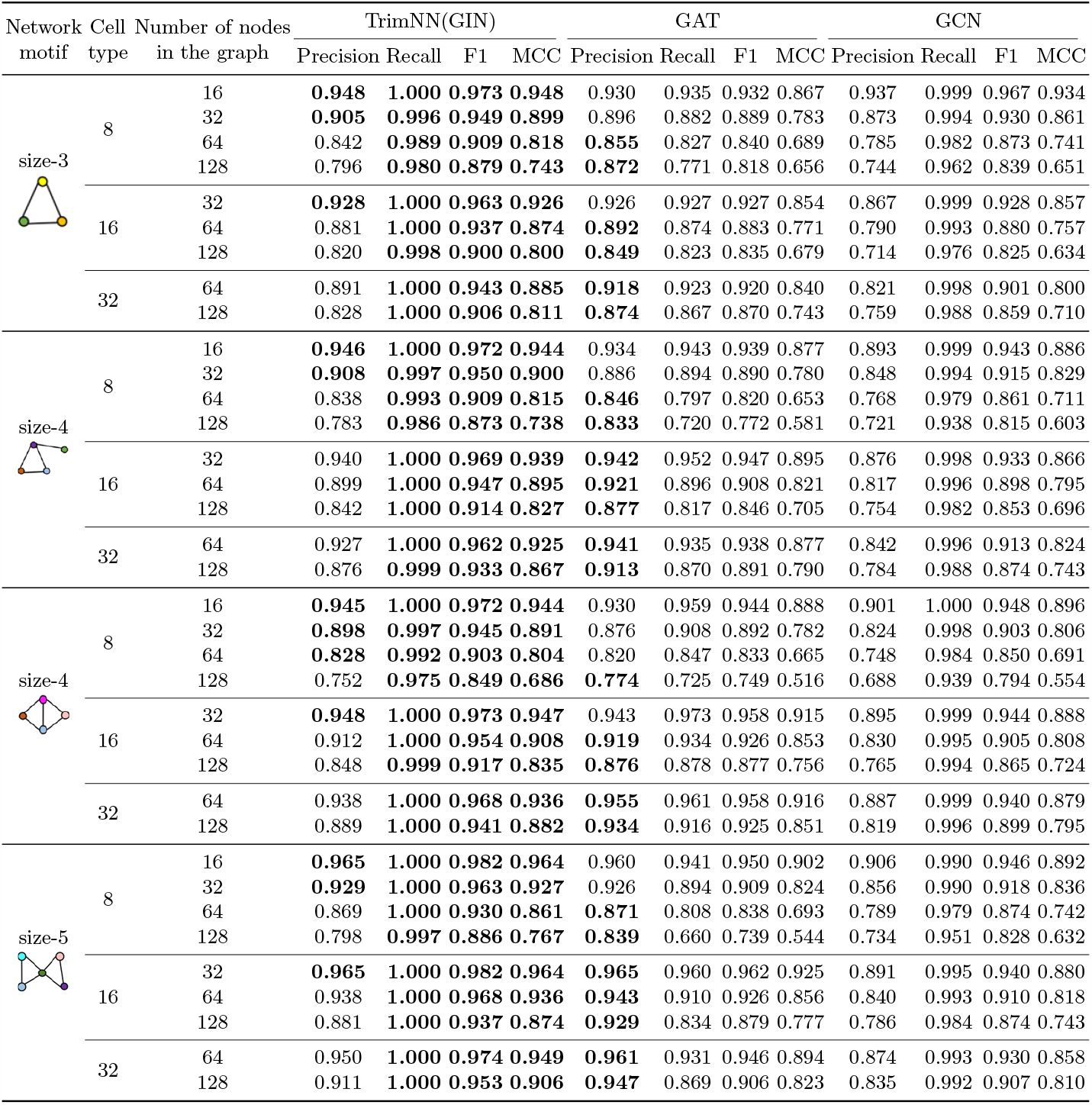
The ablation test on different graph representation methods.

| Network motif | Cell type | Number of nodes in the graph | TrimNN(GIN) |  |  |  | GAT |  |  |  | GCN |  |  |  |
| --- | --- | --- | --- | --- | --- | --- | --- | --- | --- | --- | --- | --- | --- | --- |
|  |  |  | Precision | Recall | F1 | MCC | Precision | Recall | F1 | MCC | Precision | Recall | F1 | MCC |
| size-3 | 8 | 16 | **0.948** | **1.000** | **0.973** | **0.948** | 0.930 | 0.935 | 0.932 | 0.867 | 0.937 | 0.999 | 0.967 | 0.934 |
|  |  | 32 | **0.905** | **0.996** | **0.949** | **0.899** | 0.896 | 0.882 | 0.889 | 0.783 | 0.873 | 0.994 | 0.930 | 0.861 |
|  |  | 64 | 0.842 | **0.989** | **0.909** | **0.818** | **0.855** | 0.827 | 0.840 | 0.689 | 0.785 | 0.982 | 0.873 | 0.741 |
|  |  | 128 | 0.796 | **0.980** | **0.879** | **0.743** | **0.872** | 0.771 | 0.818 | 0.656 | 0.744 | 0.962 | 0.839 | 0.651 |
|  | 16 | 32 | **0.928** | **1.000** | **0.963** | **0.926** | 0.926 | 0.927 | 0.927 | 0.854 | 0.867 | 0.999 | 0.928 | 0.857 |
|  |  | 64 | 0.881 | **1.000** | **0.937** | **0.874** | **0.892** | 0.874 | 0.883 | 0.771 | 0.790 | 0.993 | 0.880 | 0.757 |
|  |  | 128 | 0.820 | **0.998** | **0.900** | **0.800** | **0.849** | 0.823 | 0.835 | 0.679 | 0.714 | 0.976 | 0.825 | 0.634 |
|  | 32 | 64 | 0.891 | **1.000** | **0.943** | **0.885** | **0.918** | 0.923 | 0.920 | 0.840 | 0.821 | 0.998 | 0.901 | 0.800 |
|  |  | 128 | 0.828 | **1.000** | **0.906** | **0.811** | **0.874** | 0.867 | 0.870 | 0.743 | 0.759 | 0.988 | 0.859 | 0.710 |
| size-4 | 8 | 16 | **0.946** | **1.000** | **0.972** | **0.944** | 0.934 | 0.943 | 0.939 | 0.877 | 0.893 | 0.999 | 0.943 | 0.886 |
|  |  | 32 | **0.908** | **0.997** | **0.950** | **0.900** | 0.886 | 0.894 | 0.890 | 0.780 | 0.848 | 0.994 | 0.915 | 0.829 |
|  |  | 64 | 0.838 | **0.993** | **0.909** | **0.815** | **0.846** | 0.797 | 0.820 | 0.653 | 0.768 | 0.979 | 0.861 | 0.711 |
|  |  | 128 | 0.783 | **0.986** | **0.873** | **0.738** | **0.833** | 0.720 | 0.772 | 0.581 | 0.721 | 0.938 | 0.815 | 0.603 |
|  | 16 | 32 | 0.940 | **1.000** | **0.969** | **0.939** | **0.942** | 0.952 | 0.947 | 0.895 | 0.876 | 0.998 | 0.933 | 0.866 |
|  |  | 64 | 0.899 | **1.000** | **0.947** | **0.895** | **0.921** | 0.896 | 0.908 | 0.821 | 0.817 | 0.996 | 0.898 | 0.795 |
|  |  | 128 | 0.842 | **1.000** | **0.914** | **0.827** | **0.877** | 0.817 | 0.846 | 0.705 | 0.754 | 0.982 | 0.853 | 0.696 |
|  | 32 | 64 | 0.927 | **1.000** | **0.962** | **0.925** | **0.941** | 0.935 | 0.938 | 0.877 | 0.842 | 0.996 | 0.913 | 0.824 |
|  |  | 128 | 0.876 | **0.999** | **0.933** | **0.867** | **0.913** | 0.870 | 0.891 | 0.790 | 0.784 | 0.988 | 0.874 | 0.743 |
| size-4 | 8 | 16 | **0.945** | **1.000** | **0.972** | **0.944** | 0.930 | 0.959 | 0.944 | 0.888 | 0.901 | 1.000 | 0.948 | 0.896 |
|  |  | 32 | **0.898** | **0.997** | **0.945** | **0.891** | 0.876 | 0.908 | 0.892 | 0.782 | 0.824 | 0.998 | 0.903 | 0.806 |
|  |  | 64 | **0.828** | **0.992** | **0.903** | **0.804** | 0.820 | 0.847 | 0.833 | 0.665 | 0.748 | 0.984 | 0.850 | 0.691 |
|  |  | 128 | 0.752 | **0.975** | **0.849** | **0.686** | **0.774** | 0.725 | 0.749 | 0.516 | 0.688 | 0.939 | 0.794 | 0.554 |
|  | 16 | 32 | **0.948** | **1.000** | **0.973** | **0.947** | 0.943 | 0.973 | 0.958 | 0.915 | 0.895 | 0.999 | 0.944 | 0.888 |
|  |  | 64 | 0.912 | **1.000** | **0.954** | **0.908** | **0.919** | 0.934 | 0.926 | 0.853 | 0.830 | 0.995 | 0.905 | 0.808 |
|  |  | 128 | 0.848 | **0.999** | **0.917** | **0.835** | **0.876** | 0.878 | 0.877 | 0.756 | 0.765 | 0.994 | 0.865 | 0.724 |
|  | 32 | 64 | 0.938 | **1.000** | **0.968** | **0.936** | **0.955** | 0.961 | 0.958 | 0.916 | 0.887 | 0.999 | 0.940 | 0.879 |
|  |  | 128 | 0.889 | **1.000** | **0.941** | **0.882** | **0.934** | 0.916 | 0.925 | 0.851 | 0.819 | 0.996 | 0.899 | 0.795 |
| size-5 | 8 | 16 | **0.965** | **1.000** | **0.982** | **0.964** | 0.960 | 0.941 | 0.950 | 0.902 | 0.906 | 0.990 | 0.946 | 0.892 |
|  |  | 32 | **0.929** | **1.000** | **0.963** | **0.927** | 0.926 | 0.894 | 0.909 | 0.824 | 0.856 | 0.990 | 0.918 | 0.836 |
|  |  | 64 | 0.869 | **1.000** | **0.930** | **0.861** | **0.871** | 0.808 | 0.838 | 0.693 | 0.789 | 0.979 | 0.874 | 0.742 |
|  |  | 128 | 0.798 | **0.997** | **0.886** | **0.767** | **0.839** | 0.660 | 0.739 | 0.544 | 0.734 | 0.951 | 0.828 | 0.632 |
|  | 16 | 32 | **0.965** | **1.000** | **0.982** | **0.964** | **0.965** | 0.960 | 0.962 | 0.925 | 0.891 | 0.995 | 0.940 | 0.880 |
|  |  | 64 | 0.938 | **1.000** | **0.968** | **0.936** | **0.943** | 0.910 | 0.926 | 0.856 | 0.840 | 0.993 | 0.910 | 0.818 |
|  |  | 128 | 0.881 | **1.000** | **0.937** | **0.874** | **0.929** | 0.834 | 0.879 | 0.777 | 0.786 | 0.984 | 0.874 | 0.743 |
|  | 32 | 64 | 0.950 | **1.000** | **0.974** | **0.949** | **0.961** | 0.931 | 0.946 | 0.894 | 0.874 | 0.993 | 0.930 | 0.858 |
|  |  | 128 | 0.911 | **1.000** | **0.953** | **0.906** | **0.947** | 0.869 | 0.906 | 0.823 | 0.835 | 0.992 | 0.907 | 0.810 |

#### TrimNN needs sufficient training data to achieve the best performance

Another critical factor that determines the performance of TrimNN is the size of the training set. In this ablation test, we reduced the number of utilized training pairs of subgraphs and triangulated graphs to half and one-fourth, then retrained the model. The model performance are presented in Table 4 on dataset and criteria in Section 3.1. We can see that as the size of the training dataset decreases, the model’s performance deteriorates quickly. This phenomenon indicates that TrimNN model requires sufficient training data to effectively learn the complex present/absent relationships between subgraphs and triangulated graphs.

**Table 4.**
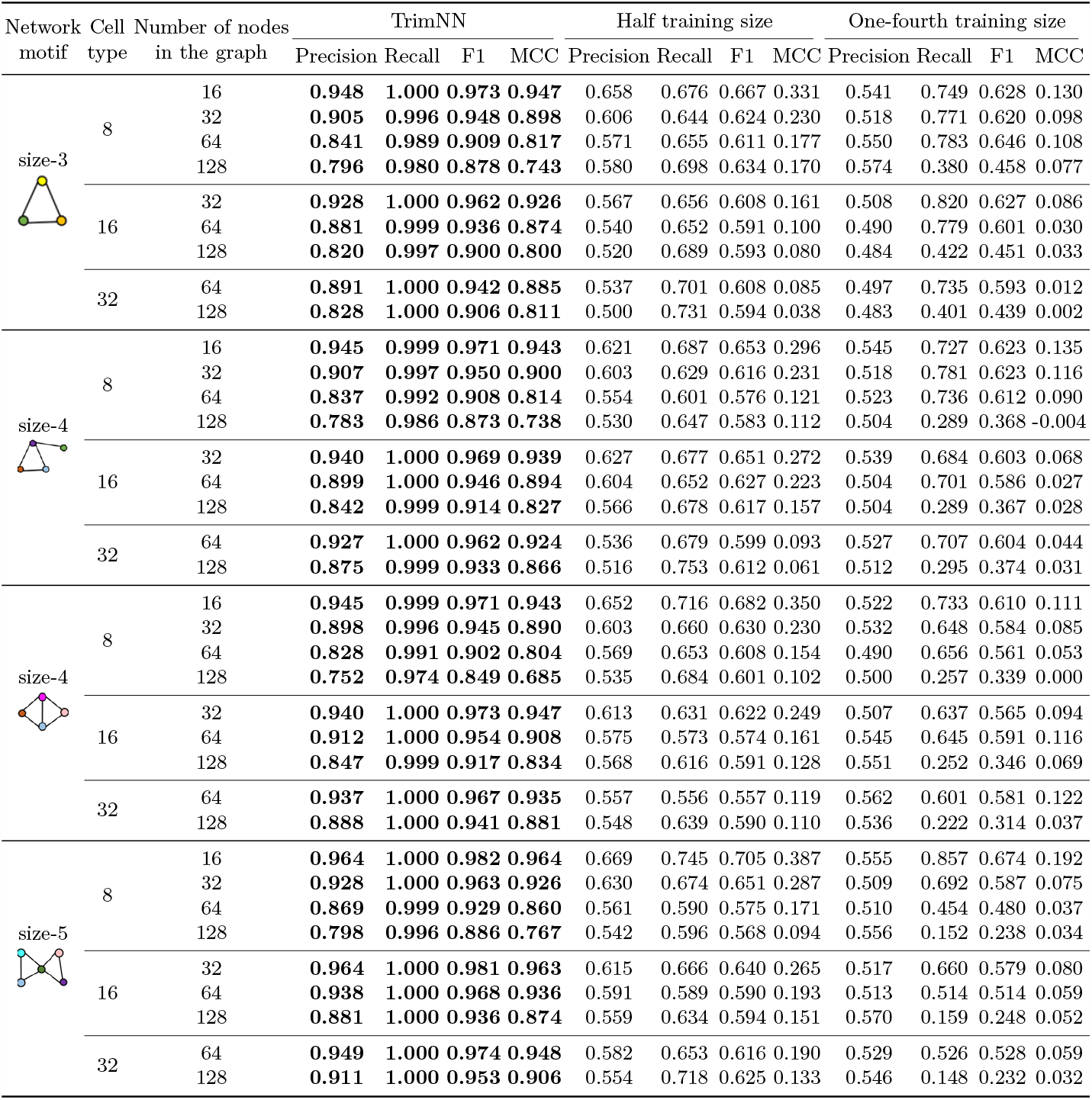
The major performance of different training data sizes.

### 3.5 Case study on Alzheimer’s disease

We analyzed brain tissues of a mouse model of Alzheimer’s disease (AD) at 8 months of age. These SRT data are generated with STARmap PLUS[30] technologies at subcellular resolution. The nodes’ cell types are annotated by the original publication. We systematically screened size-4 network motifs on case and control samples. Among them, a network motif with cell type cortex excitatory neuron, microglia, and dentate gyrus is found to be overrepresented in AD samples with 251 occurrences. However, it only occurred 3 times in the control sample with an odds ratio of ∼83.7. These preliminary results may be interpreted as microglia and cortex’s unique transcriptional changes and interplay in AD[17]. This network motif is visualized in **Supplementary Figure 1**.

## 4 Discussion

The advent of spatial omics has revolutionized our capacity to explore the nuanced organization of cells within tissues at the cellular level. Based on graph isomorphism network, the proposed work formulates the pattern quantification problem in counting subgraph occurrences, simplifying the task to a biologically constrained problem. TrimNN provides an accurate, unbiased, efficient, and robust approach to quantify the network motifs as interpretable building blocks of cell organization. The success of TrimNN relies on 1) simplifying the NP-complete subgraph matching problem on the universal graph to a well-defined set of biological meaningful triangulated graphs, 2) decomposing the challenging isomorphic counting regression problem on the entire graph to many straightforward binary present/absent prediction problems on small graphs, and 3) using supervised learning to achieve scalability in inferences. This work paves the way to disclose the biological mechanisms underlying multicellular differentiation, development, and disease progression.

There are still some limitations of this work. Firstly, TrimNN mainly contributes to dealing with the subgraph matching problem with a given subgraph, we still need to enumerate or search numerous subgraphs to get the top ones. Then, the results from TrimNN are still estimated, with no guarantee of exact results like the enumeration methods. In the future, the identified enriched network motifs in different categories of diseases will be evaluated with downstream analysis, such as cell-cell communication annotation[6], clinical information, and other biological validations[13] from independent data sources.

## Notes

### Competing Interest Statement

The authors have declared no competing interest.

